# The microRNomes of Chinese Hamster Ovary (CHO) cells and their extracellular vesicles, and how they respond to osmotic and ammonia stress

**DOI:** 10.1101/2022.12.16.520798

**Authors:** Jessica Belliveau, Eleftherios T. Papoutsakis

**Affiliations:** Department of Chemical and Biomolecular Engineering, Newark, DE 19711; Delaware Biotechnology Institute, Newark, DE 19711; Department of Biological Sciences, University of Delaware, Newark, DE 19711

**Keywords:** Chinese hamster ovary cells, extracellular vesicles, miR, ammonia stress, osmotic stress

## Abstract

A new area of focus in Chinese Hamster Ovary (CHO) biotechnology is the role of small (exosomes) and large (microvesicles or microparticles) extracellular vesicles (EVs). CHO cells in culture exchange large quantities of proteins and RNA through these EVs, yet the content and role of these EVs remain elusive. MicroRNAs (miRs) are central to adaptive responses to stress and more broadly to changes in culture conditions. Given that EVs are highly enriched in miRs, and that EVs release large quantities of miRs both *in vivo* and *in vitro*, EVs and their miR content likely play an important role in adaptive responses. Here we report the miRNA landscape of CHO cells and their EVs under normal culture conditions and under ammonia and osmotic stress. We show that both cells and EVs are highly enriched in five miRs (among over 600 miRs) that make up about half of their total miR content, and that these highly enriched miRs differ significantly between normal and stress culture conditions. Notable is the high enrichment in mir-92a and miR-23a under normal culture conditions, in contrast to the high enrichment in let-7 family miRs (let-7c, let-7b and let-7a) under both stress conditions. The latter suggests a preserved stress-responsive function of the let-7 miR family, one of the most highly preserved miR families across species, where among other functions, let-7 miRs regulate core oncogenes, which, depending on the biological context, may tip the balance between cell cycle arrest and apoptosis. While the expected –based on their profound enrichment – important role of these highly enriched miRs remains to be dissected, our data and analysis constitute an important resource for exploring the role of miRs in cell adaptation as well as for synthetic applications.

## 1. INTRODUCTION

Chinese hamster ovary (CHO) cells are the dominant mammalian host cell line for the production of recombinant biotherapeutics. CHO culture performance and optimization is a key goal in CHO biotechnology. While the sequence of the CHO genome has been largely completed (e.g., (Hilliard, MacDonald, & Lee, 2020; Rupp et al., 2018)), its annotation and functional behavior remain to a large extent not well understood. As each producer cell line is different from all others due to selection and cloning, global traits of how cells respond to environmental changes – important in process optimization – are likely preserved. Cells respond to environmental conditions by altering their transcriptional program as a first line of global adaptation. Significantly, small RNAs, and notably miRNAs (miRs), play an important role in fast adaptation and responses to environmental changes and/or stress as miRs bridge the transcriptional to translational nexus (Jadhav et al., 2013). Yet their role remains ill understood due to their highly multiplexed mechanisms of action. Understanding adaptational responses may enable diagnostic and synthetic applications in cell engineering and will illuminate cell physiology (Jadhav et al., 2013; Stiefel et al., 2015). While several studies have examined the total CHO RNome of normal cultures and strains using RNASeq (e.g., (Bort et al., 2012; C. Chen, Le, & Goudar, 2017; K. M. Chen et al., 2017; Fomina-Yadlin et al., 2015; Konitzer et al., 2015; Orellana et al., 2018; Sha, Bhatia, & Yoon, 2018; Vishwanathan et al., 2015)), transcriptional responses to physiological stressors remain largely unexplored (Synoground et al., 2021).

MicroRNA (miRNA or miR) are small, non-coding RNAs averaging 22 nucleotides in length that are involved in a multitude of regulatory mechanisms. miRs transcribed in the nucleus form primary miRNAs (pri-miRNAs), which are cleaved at the base of the hairpin structure to form precursor miRNAs (pre-miRNAs) and are exported into the cytoplasm. Pre-miRNA is cleaved at the top loop structure, resulting in two mature miRNAs, the 5p strand that originated from the 5’ end of the pre-miRNA and the 3p strand that originated from the 3’ end, and can both regulate gene expression (Hackl et al. (2011); O’Brien, Breyne, Ughetto, Laurent, and Breakefield (2020)). The Argonaute (AGO) protein binds mature miRNAs to form the RNA-induced silencing complex (RISC) that can silence genes with complementary sequence motifs to the loaded miRNA.

Extracellular vesicles (EVs) are produced by all cells and function as communication messengers by transporting cargo of proteins, RNA, and lipids between cells, both in vivo and in vitro (C.-Y. Kao and Papoutsakis (2018); C. Y. Kao and Papoutsakis (2019); Raposo and Stoorvogel (2013)). Microparticles (MPs) – also referred to as microvesicles (MVs) – are a class of EVs formed from the budding of the cell membrane and are typically between 100 nm and 1 μm. Exosomes, the second primary class of EVs, are formed from the endosomal membrane and are typically less than 150 nm in diameter (Jiang, Kao, and Papoutsakis (2017); Raposo and Stoorvogel (2013)). EV-mediated transfer of RNA and other cargo to target cells is not well understood mechanistically, but can affect the target cells in a multitude of ways (Adamo, Dal Collo, Bazzoni, & Krampera, 2019; Chin & Wang, 2016; de Andrade, Mehta, & Bresnick, 2020; Jiang et al., 2017; C. Y. Kao, Jiang, Thompson, & Papoutsakis, 2022; C. Y. Kao & Papoutsakis, 2019; Sandiford et al., 2021). In terms of relative abundance, EVs are enriched in small RNAs (miRNA, siRNA, lncRNA, piRNA) compared to the parent cell (Corrado, Barreca, Zichittella, Alessandro, and Conigliaro (2021)). While several studies have reported changes in cell phenotype and expression when specific EVs are cocultured with various targets (Montecalvo et al. (2012); M. Yang et al. (2011)), other studies suggest the quantity of these small RNAs may not be sufficient to alter the cellular state (O’Brien et al. (2020)). Changes in the EV small RNA profile or protein content may indicate changes in the cellular phenotype and be used as a culture diagnostic tool (Kosaka et al. (2019)).

The RNA profile of EVs is not a direct representation of the parent cell (O’Grady et al. (2022); Sherman, Lodha, and Sahoo (2021)). Cargo loading of EVs is apparently a selective process where RNAs with specific motifs are actively enriched (Corrado et al. (2021)). These loading mechanisms may differ for microparticles and exosomes and among cell types, and overall are not well understood (Corrado et al., 2021). We have demonstrated in in vitro cultures of CHO cells and human primary and immortalized cells that there is a massive, dynamic process whereby EVs are continuously produced and taken up by cells (Belliveau and Papoutsakis (2022)) leading to widespread exchange of proteins and RNAs via EVs between the cells in the same culture. In view of the extensive exchange of RNAs between cells in culture through EVs, and the central role of miRs in the cellular adaptive responses to stress (Anthony K. L. Leung & Sharp, 2007; A. K. L. Leung & Sharp, 2010)), here, we aim to explore the microRNA landscape of CHO cells and their EVs in standard cultures and in cultures under two stress conditions, ammonia and osmolarity.

CHO miRs have been examined in a few studies over the last 15 years (Bort et al., 2012; Busch et al., 2022; Hackl et al., 2014; Jadhav et al., 2013; Keysberg et al., 2021; Klanert et al., 2016; Maccani et al., 2014) and have been also used in cell engineering applications (e.g., (Druz, Son, Betenbaugh, & Shiloach, 2013; Jadhav et al., 2013; Jadhav et al., 2014; Raab et al., 2019). Here, given the profound exchange of miRs between cells through EVs, we aimed to examine how the miRNA content of cells and their EVs compare, how does that content changes with stress, and what are the most abundant miRs under standard and stress culture conditions. The current literature does not provide a clear picture of what are the most abundant miRs in CHO cells and what is their relative abundance under normal or stress culture conditions. Such information would be important in functional miR assignment if one makes the hypotheses that the most abundant miRs in cells and perhaps EVs play a role in cell proliferation and metabolism, and that changes in the most abundant miRs would reflect the impact of these miRs on cellular adaptation to altered environmental conditions. While not specifically examined for CHO cells or stressors like ammonia and high osmotic pressure, as recently reviewed (Olejniczak, Kotowska-Zimmer, & Krzyzosiak, 2018), experimental data support the notion that cellular stress directly influences not only the miRNA concentration but also the function miR biogenesis proteins and the final sequence of mature miRs.

## 2. MATERIALS AND METHODS

### 2.1 Cell-culture setup

A recombinant CHO-K1 (Clone A11 from the Vaccine Research Center at the National Institutes of Health) cell line expressing an anti-HIV VRC01 antibody was cultured in HyClone ActiPro media (Cytiva, Marlborough, MA) supplemented with 6 mM L-glutamine in 50 mL conical culture tubes at 250 RPM at 37°C in 5% CO_2_. Cultures were seeded at a density of 0.4x10^6^ cells/mL in a total volume of 15 mL. Ammonium chloride (NH_4_Cl) dissolved in PBS (pH 7.4), was added to ammonia stressed cultures at the start of culture for a starting concentration of 9 mM NH_4_Cl. Sodium chloride (NaCl) dissolved in PBS (pH 7.4), was added to osmotic stressed cultures at the start of culture, corresponding to an additional 120 mOsm/L, for a starting concentration of 60 mM NaCl. All cultures for analysis (cell concentration, viability, EV size distribution, RNA sequencing) were grown with three biological replicates.

### 2.2 Isolation and characterization of extracellular vesicles

EVs were isolated with differential ultracentrifugation following the protocol previously described (Belliveau and Papoutsakis (2022)). Briefly, cells were first removed from the culture by centrifugation at 180*g* for 5 minutes, then cellular debris and apoptotic bodies removed by centrifugation at 2000*g* for 10 minutes. The supernatant was centrifuged at 28,000*g* for 30 minutes (Beckman Coulter Optima LE-80K Ultracentrifugation, SW-28 rotor) and concentrated at 28,000*g* for 30 minutes with the Beckman Coulter Optima MAX Ultracentrifuge (TLA-55 rotor) to isolate MPs. Isolated MPs were kept in growth media and used immediately or stored at 4°C overnight for use the following day. After the MPs were removed, the culture supernatant was passed through a 0.22 μm filter to remove any remaining MPs and centrifuged at 100,000*g* for 90 minutes (SW-28 rotor, Beckman Coulter Optima LE-80K). Exosomes were concentrated with the Beckman Coulter Optima MAX Ultracentrifuge (TLA-55 rotor) at 100,000*g* for 90 minutes. Isolated exosomes were kept in growth media and used immediately or stored at 4°C overnight for use the following day. Microparticle count and concentration was determined by flow cytometry using the Violet Side Scatter detector on the CytoFlex S (Beckman Coulter) flow cytometer.

### 2.3 Nanoparticle Tracking Analysis (NTA)

EVs were counted with Nanoparticle Tracking Analysis (Nano-Sight NS3000, Malvern Panalytical) at a 100-fold dilution in filtered PBS. The size distribution for each EV condition was averaged between three biological replicates.

### 2.4 Transmission electron microscopy (TEM) analysis

Isolated EVs were placed on 400 mesh carbon coated copper grides (Electron Microscopy Sciences) and stained with uranyl acetate for negative staining for transmission electron microscopy (TEM) imaging (Libra 120, Zeiss)

### 2.5 RNA sequencing (RNASeq)

For standard exponential phase cultures, CHO cultures (3 biological replicates) were grown under standard culture conditions for 3 days before cells, MPs, and exosomes were harvested as described above. For stressed cultures, CHO cultures (3 biological replicates) were grown as described above and harvested on day 3 of culture. Total RNA was harvested from cells, MPs and exosomes with the RNeasy mini kit (Qiagen) and eluted from the column in RNase, DNase free water. A 5 uL aliquot of the RNA was used for fragment analysis (AATI Fragment Analyzer, Agilent) to determine RNA size distribution, quality, and quantification. RNA concentration was determined by Qubit RNA HS Assay kit (ThermoFisher). RNA sequencing was completed in two rounds, first with the samples from standard cultures and second with the samples from stressed cultures, using two different library preparation kits. Total RNA from standard cultures were prepared according to manufacturer’s protocol using the NextFlex Combo-Seq mRNA/miRNA kit (PerkinElmer), to capture both the mRNA and miRNA in the samples. Due to low mRNA read counts with the Combo-Seq kit, we focused solely on the small RNA, therefore, the total RNA from stressed cultures was prepared using the TruSeq Small RNA Sample Prep Kit (Illumina). Sequencing of was done on one run of Illumina NextSeq 550 producing ∼360M single-end 76 bp reads (range per sample: 32.6-45.8M). Library preparation and RNA sequencing was done through the UD Sequencing & Genotyping Center and the data analysis was performed by the UD Bioinformatics & Computational Biology Core.

### 2.6 Analysis of RNA sequencing data

Sequencing data analysis was provided by Dr. Shawn Polson and Jaysheel Bhavsar (Center for Bioinformatics and Computational Biology) using the established RNA-Seq analysis pipeline (adapted from Kalari et al., 2014). Quality of sequencing data was assessed using FastQC (ver. 0.10.1; Babraham Bioinformatics). All reads were mapped to the Cricetulus griseus genome PICR assembly (RefSeq GCF_003668045.1, Annotation version 103)14 to determine the overall biotype (e.g. mRNA, miRNA, etc.) composition of reads using CLC Genomics Map Reads to Reference tool with default settings (ver 20.0.2).

Candidate small RNA sequences were identified from raw sequence data using cutadapt (ver. 2.8)21 with parameters “-u -4 -a A(10)” as recommended by PerkinElmer NextFlex Combo-Seq kit. All reads shorter than 50 bp and containing the poly-A adapter were utilized for miRNA and piRNA analysis. Reads passing filter were analyzed using the CLC Genomics Server (ver. 20.0.2; Qiagen Bioinformatics). Samples were processed in batch using CLC Quantify miRNA with “strand specific=yes” and allowing up to 2 mismatches. For miRNA, miRbase (ver. 22.1)22 was used as a reference database with species in this priority order: C. griseus, Mus musculus, Rattus norvegicus, and Homo sapiens and results were grouped on mature miRNA exact matches. For piRNA the piRNAdb for C. griseus (cgr.v1.7.6; https://www.pirnadb.org) was used as reference databases and counts were grouped on 5’ mature matches including variants. Differential expression analysis utilized TMM normalization and the CLC Differential Expression Analysis based on negative binomial Generalized Linear Model (GLM) using the Wald test for determining statistical significance23 and FDR correction24.

## 3. RESULTS

### 3.1 Characterization of CHO stressed and standard cultures

Two stress conditions were selected for investigation of changes to the microRNome of cells and EVs: ammonia stress and osmolarity stress. All stressors were added at the start of the culture. We chose to identify stress levels that reduced cell growth but not cell viability. Lactate stress was also investigated, however due to no changes in cell growth compared to the control culture at a range of 30 mM to 120 mM of added sodium lactate (data not shown), lactate stress was not further pursued. In selecting an appropriate stress, three concentrations of NH_4_Cl (3 mM, 6 mM, and 9 mM) for ammonia stress or sodium chloride (30 mM, 60 mM, and 120 mM) for osmotic stress were tested based on literature values (Kiehl, Shen, Khattak, Jian Li, & Sharfstein, 2011; Lao & Toth, 1997; Synoground et al., 2021). Changes in cell concentration was the preliminary screen for determining the appropriate stressor level. Significant differences in cell concentrations (Figure 1a) for ammonia and osmotically stressed cultures compared to the standard cultures were observed at days 1 to 4. After four days of culture, the cell concentration of cultures with 9 mM NH_4_Cl stress was 59% of the standard culture and with an additional 120 mOsm/L added stress (for osmotic stress) was 43% of the standard culture. Stresses impacted the overall culture growth, but the culture was still able to maintain exponential growth. Cell viability (Figure 1b) was measured over the four days of culture with Trypan Blue staining.

**Figure 1:**
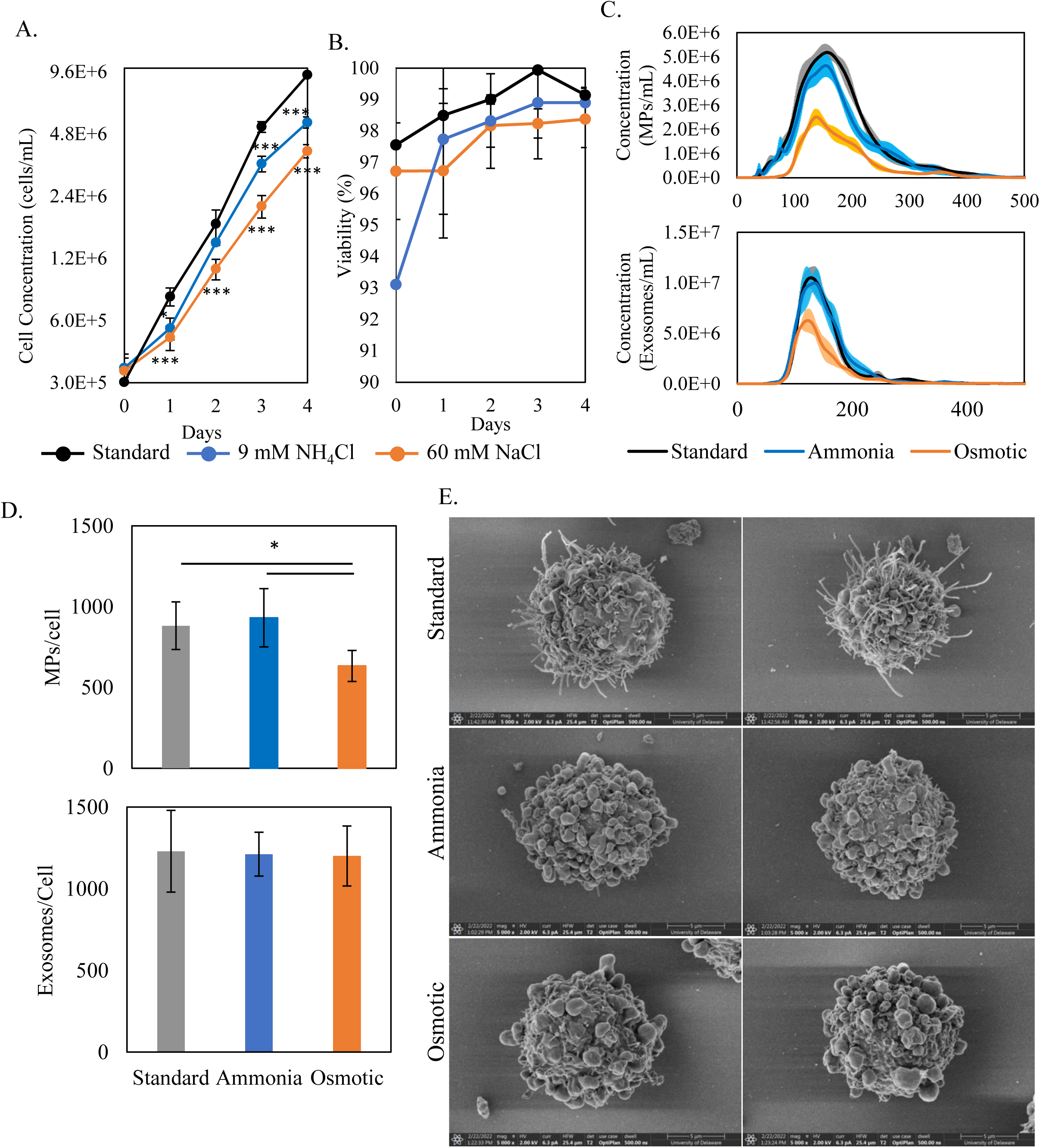
Characterization of stressed and unstressed cultures. A. Cell growth curve of cultures growth with an added 9 mM ammonia chloride for ammonia stressed cultures (blue), 60 mM sodium chloride for osmotic stressed cultures (orange), and standard, no added stress cultures (black). B. Cell viability measured with Trypan Blue staining of ammonia, osmotic, and standard cultures. C. NTA measurements of microparticle (upper) and exosome (lower) size distribution. D. Calculation of microparticles and exosomes per cell in culture. E. Representative SEM images of cells from standard, ammonia, or osmotic stressed cultures. Three biological replicates, student t-test with p-values of * 0.05, **0.01, and ***<0.001.

Viability was greater than 96% for both standard and stressed cultures. Although stressed cultures had somewhat lower viabilities, differences were not statistically significant. Thus, while there were significant differences in cell concentrations, there were no significant differences in cell viability, indicating an appropriate stress level to further explore.

Microparticles and exosomes were isolated from the stressed and standard cultures on day 3 via differential ultracentrifugation and filtration for size distribution and concentration analysis via nanoparticle tracking analysis (NTA) (Figure 1c). The size distributions of the MPs and exosomes from the ammonia stressed cultures were similar to those of the standard cultures with the respective size distribution peaks of 154 nm and 157 nm for the MPs and 134 nm and 128 nm for the exosomes. In Figure 1d, when EV concentrations were normalized to cell concentration, there was no difference in the number MPs or exosomes produced by ammonia stressed cells compared to standard cultures. MPs from osmotically stressed cultures were smaller than the standard cultures with the MP size distribution peak at 138 nm (Figure 1c), however the peak of the exosomes, 122 nm, was similar to the standard cultures. The normalized number of MPs produced per osmotically stressed cell was significantly lower than the number of MPs produced per cell in standard or and ammonia stressed cultures. However, there was no differences in the number of exosomes produced per cell. The differences in EV production per cell due to ammonia or osmotic stress conditions suggests that stresses affect EV production differently and that MP and exosome production per cell are independently affected by these stresses.

The morphology of cells in stressed cultures was markedly different from cells in standard cultures. As previously presented (Belliveau & Papoutsakis, 2022), in exponential phase of standard cultures, cells exhibit long microvilli that cover a large portion of the cell surface and EV-like structures attached to the cell as shown in two representative SEM images in Figure 1e. Cells from ammonia and osmotic stressed cultures lack the long microvilli structure and the cell surface appear to be covered in EV-like structure. Microvilli are likely important in the ability of cells to capture and integrate EVs (Belliveau & Papoutsakis, 2022; Jiang et al., 2017). The lack of microvilli on stressed cells might suggest a reduced cellular capacity to capture and integrate EVs from the surrounding medium. If so, the concentrations of EVs under stress conditions would not be directly comparable to those of standard conditions due to reduced EV uptake by the cells. Given that EVs are quite stable in in vitro cultures, as they have half-lives of 30 min to a few hours in vivo, where they are cleared by phagocytic cells (Lai et al., 2014), the implication would then be that under these stress conditions, cells exchange less cellular material through EVs.

### 3.2 The microRNome of standard cultures

RNA sequencing of the miRs isolated from cells, MPs, and exosomes identified unique miRs, the most abundant miRs, and differentially expressed miRs. Cells, MPs, and exosomes from standard cultures share the majority of the same miRs, 439 commonly detected miRs, and there are similar numbers of unique miRs detected in each population (Figure 2a). The miRs present in MPs and exosomes is not the same as the parent cell. There were 113 unique miRs detected in both MPs and exosomes. Since all miRs in EVs are derived from the parent cell, the interpretation here should be that the unique miRs in EVs had a very low abundance in the parent cells and could not thus be picked up by RNAseq. As discussed in the introduction, it is well established in the literature that there is a selective process of cellular RNA loading to EVs, but the cargo sorting mechanism remains largely unexplored.

**Figure 2:**
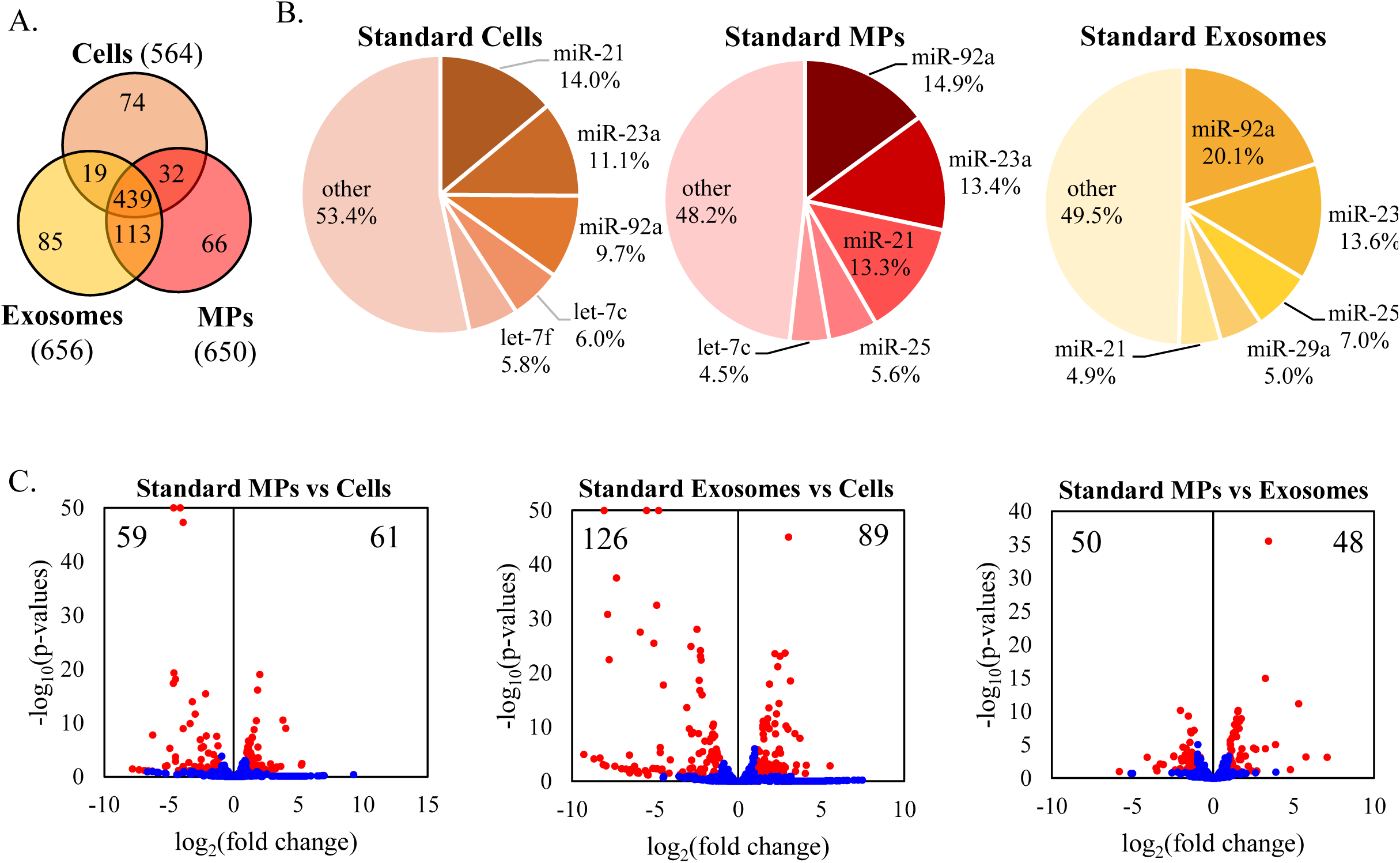
The microRNome of standard cultures. A. Venn diagram detailing the number of unique and shared miRs amongst cells, MPs, and exosomes from standard cultures. B. Pie chart of the relative abundance of the five most abundant miRs in cells, MPs, and exosomes from standard cultures. The five most abundant miRs account for approximately 50% of the total miRs detected. C. Volcano plots of the differential expression analysis of miRs with significance determined by log_2_(fold change)>1 or <1 and -log_10_(FDR p-value)<0.1. MiR profiles are the average of three biological replicates.

The relative abundance of specific miRs is shown in Figure 2b, where, among all standard cells, MPs, and exosomes, the five most abundant miRs were approximately 50% of the total miRs detected. In cells, miR-21 accounted for 14% of the total detected miRs and in MPs for 13.3%, but only 4.9% in exosomes. miR-92a abundance in cells, MPs, and exosomes was 9.7%, 14.9%, and 20.1%, respectively. In the case of miR-23a, abundance was similar across the cells, MPs, and exosomes. Roles for these three miRs, miR-21, miR-92a, and miR-23a, have been explored in human cell lines as relating to cell growth and cancer cells (Table 1), but the role these miRs as individuals or as a group have not yet been widely explored in CHO cells. miR-92 is the last miR of the well-known miR 17/92 cluster (Jadhav et al., 2013; Mogilyansky & Rigoutsos, 2013) that controls cell cycle, apoptosis, and proliferation. Assuming that this abundance/enrichment is biologically purposeful, and based on the role of these miRs in the literature (Table 1), we hypothesize that miR-21, miR-92a, and miR-23a either as individuals or as a group impact cell growth and viability and that MPs and exosomes derived from these cultures support cell growth and viability in target cells. The distribution of differentially expressed miRs (Figure 2c) in standard cultures are approximately symmetrical in the differential abundance of miR distribution. Of note, there are the large differences in number of differentially abundant miRs between cells and exosomes. Details of the specific miRs differential abundances can be found in Supplementary Table 1.

**Table 1.**
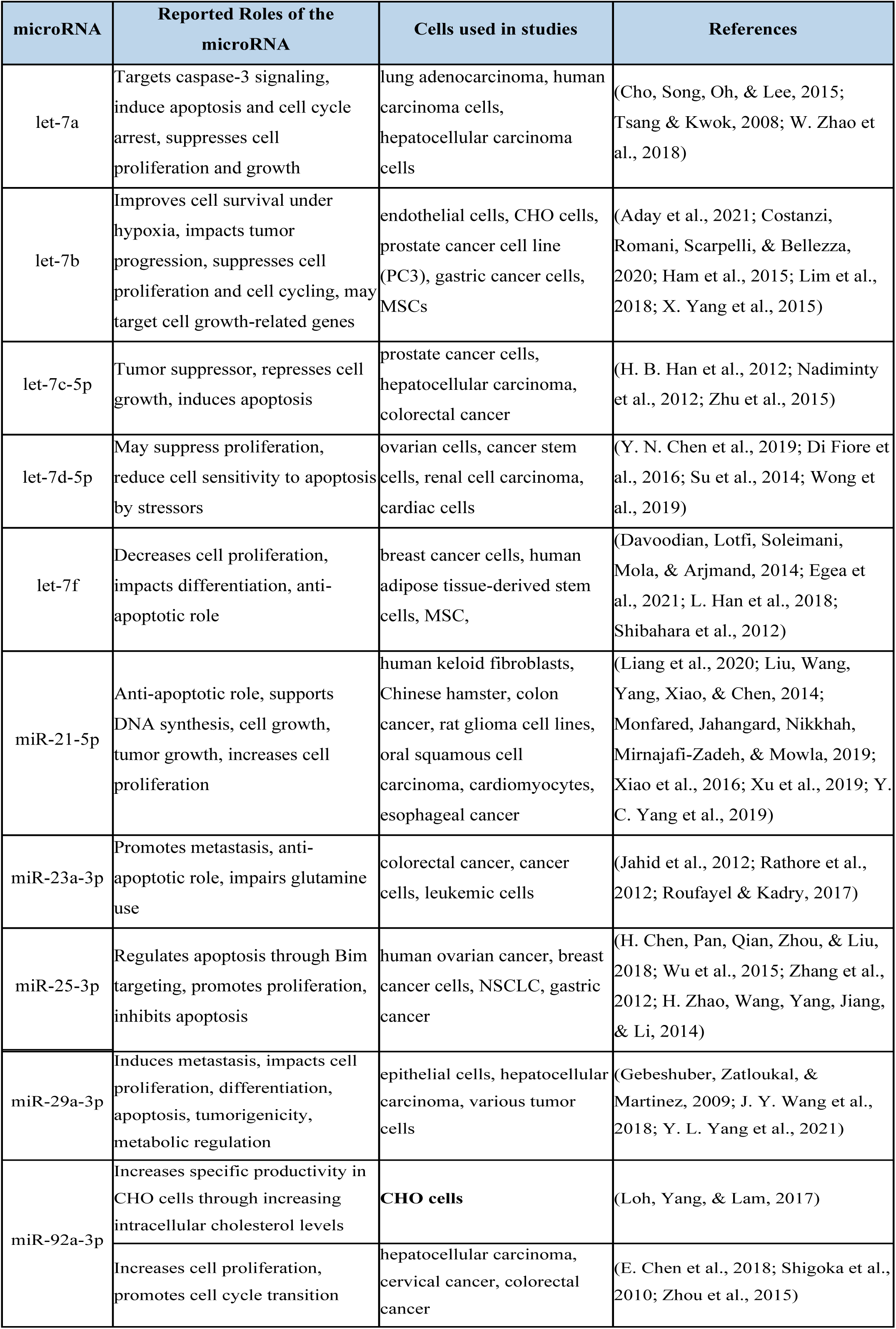

To sum, although the profile of the highly enriched miRs were similar, there were significant differences in miR content and abundances when comparing any pair of CHO cells, MPs, and exosomes, thus suggesting that although the sorting mechanisms for MPs and exosomes may be similar, the outcomes, in terms of miR cargo composition, are different.

### 3.3 The microRNome of osmotically stressed cultures

Osmotically stressed cultures resulted in a dramatic shift in the distribution of unique and shared miRs (Figure 3a) compared to the standard cultures (Figure 2a). MPs and exosomes from osmotically stressed cultures contained few unique miRs, 10 and 5 respectively, and there were no detected miRs that were found only in the MPs and exosomes. MPs and exosomes contain far fewer miRs than their counterparts of standard cultures, thus suggesting damage or inhibition of the cellular machinery that loads miRs to EVs. There were 293 unique miRs detected in osmotically stressed cells only, where the total number of miRs was similar to that of standard cultures. Of the miRs detected in MPs, 96% are shared with the parent cells. Exosomes share 98% with the parent cells.

**Figure 3:**
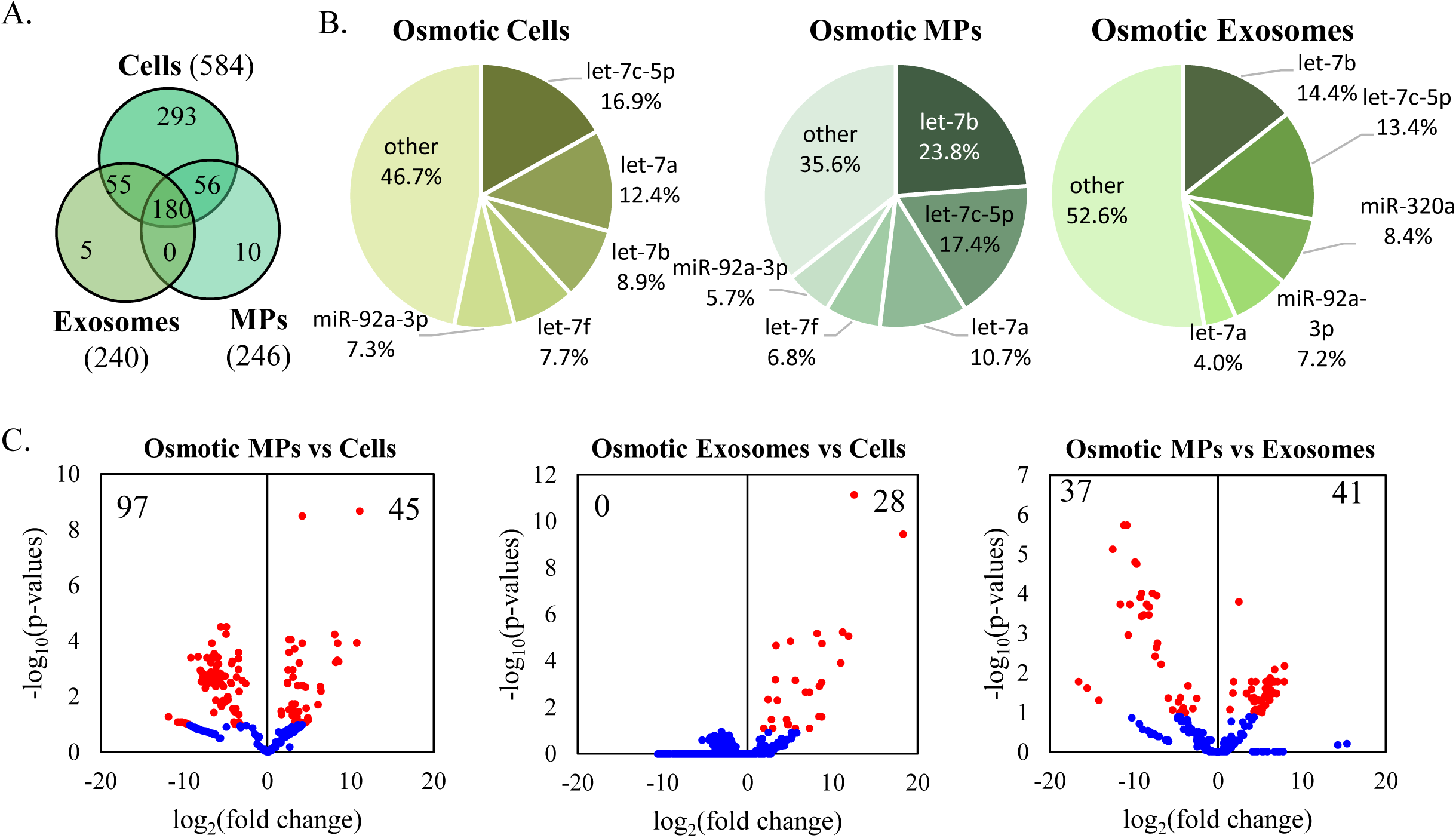
The microRNome of osmotic stressed cultures. A. Venn diagram detailing the number of unique and shared miRs amongst cells, MPs, and exosomes from osmotic stressed cultures. B. Pie chart of the relative abundance of the five most abundant miRs in cells, MPs, and exosomes from osmotic stressed cultures. The five most abundant miRs account for approximately 50% of the total miRs detected. C. Volcano plots of the differential expression analysis of miRs with significance determined by log_2_(fold change)>1 or <1 and -log_10_(FDR p-value)<0.1. MiR profiles are the average of three biological replicates.

The profile of the five most abundant miRs (Figure 3b) for the osmotically stressed cells, MPs, and exosomes largely consists of the let7 miR family. In cells and MPs, let-7a, -7b, and -7c comprise 38.2% and 51.9%, respectively of the total normalized miR counts. Highly enriched in standard cultures, miR-92a is also abundant in osmotically stressed cultures at 7.3%, 5.7%, and 7.2% of the cells, MPs, and exosomes. The miR abundance profile of the exosomes had lower levels of let-7a compared to the cells and MPs, and higher levels of miR-320a.

The role of the let7 family appears to be multifaceted and contrasting, likely depending on the let7 family member and the cell type it’s investigated in (Table 1). Let-7c and -7g have been reported to negatively regulate the anti-apoptotic oncogenes of the BCL2 family, promoting DNA damage-induced apoptosis in human hepatocellular carcinoma cells (Olejniczak et al., 2018; Shimizu et al., 2010). In human mesenchymal stem cells exposed to reactive oxygen species, let-7b has been reported to target caspase-3, suppressing apoptosis and improving cell survival (Ham et al., 2015).

Approximately twice as many miRs had differentially lower or higher abundance in osmotically stressed cells (97 miRs) compared to MPs (45 miRs) (Figure 3c). Comparing the differential miR abundance in cells vs. exosomes, there were no significantly more abundant miRs in the cells and 28 miRs show significantly lower abundance. The distribution of higher and lower in abundance miRs is approximately even when comparing MPs and exosomes, though the distribution on the volcano plot is very asymmetrical, unlike the equivalent plots from standard cultures.

Osmotic stress activates p53 (Kishi et al., 2001; Lambert, Enghoff, Brandi, & Hoffmann, 2015), the guardian of the genome, that controls cell-cycle (proliferation), apoptosis, and cycle arrest, as a means to prevent catastrophic cellular collapse. p53 regulates a large number of miRs directly or through regulation of pri-miR processing (Olejniczak et al., 2018), and that list includes the let-7 family of miRs (increased expression) and mir-17-92 cluster (reduced expression), a pattern that fits with our results here. However, p53 has been found to also upregulate mir-23a and positively control the processing of pre-miR-21, which may appear opposite to the observed pattern (Table 2). However, the relative abundance data we show do not correlate to absolute increase or decrease levels in the cell or EVs, and thus further data analysis and additional experimentation are needed. Nevertheless, it should not escape attention to the fact that virtually all the top most abundant miRs are under the control of p53 (Olejniczak et al., 2018).

**Table 2.**
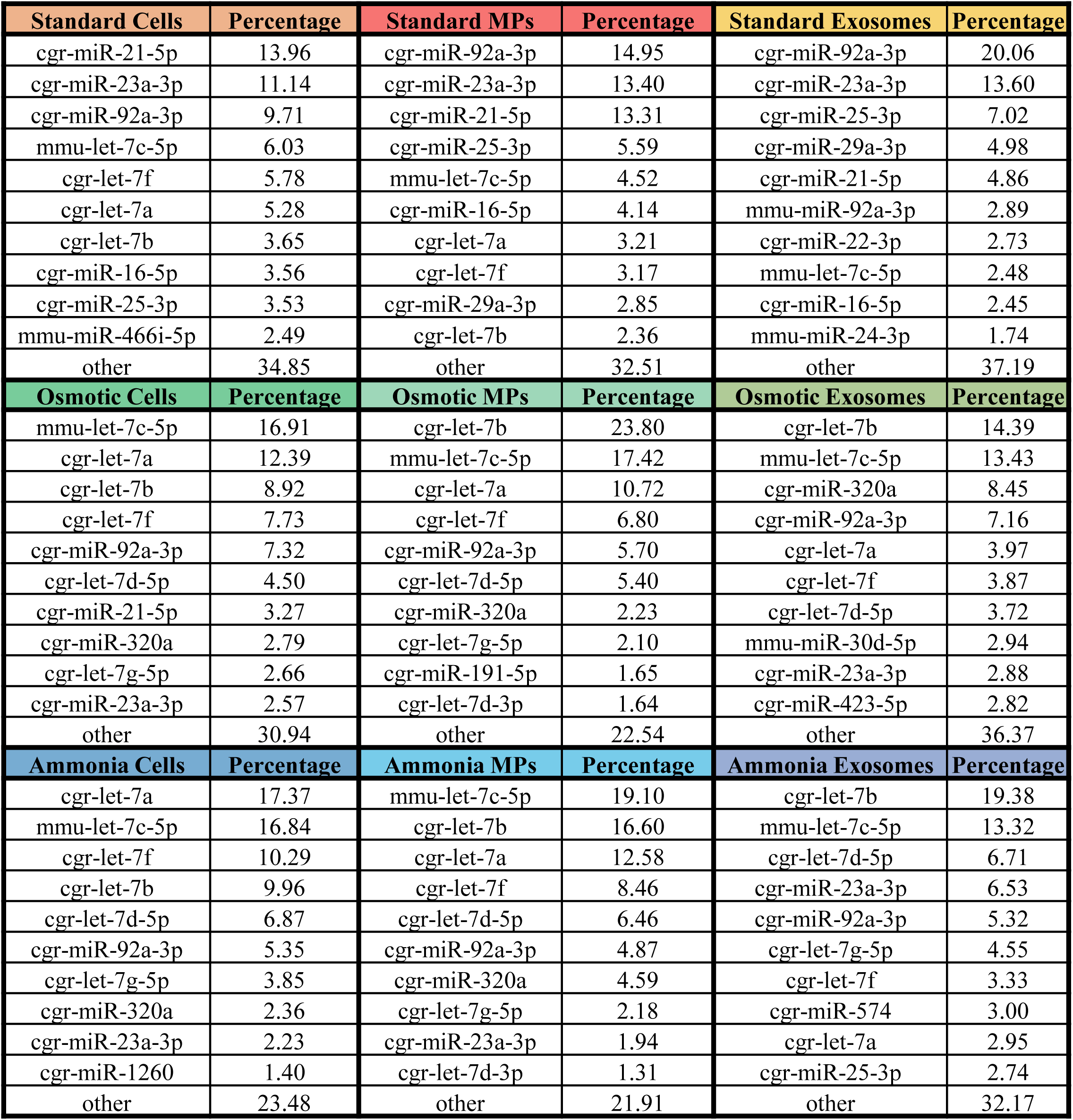

### 3.4 The microRNome of ammonia stressed cultures

In ammonia stressed cultures, MPs and exosomes contained few unique miRs (Figure 4a) compared to the parent cells and to EVs from standard cultures. Both MPs and exosomes contained >50% fewer miRs compared to their counterparts from standard cultures, again suggesting damage or inhibition of the EV sorting and loading machinery. Of the miRs detected in MPs, 93% are shared with the parent cells and exosomes share 96% with the parent cells. Like in the osmotically stressed cultures, the let7 family of miRs are highly enriched in cells, MPs, and exosomes from ammonia stressed cultures (Figure 4b). Similar to the situation of osmotically stress cultures, the five most abundant miRs in ammonia stressed cells and MPs are in the let-7 family, let-7a, -7b, -7c, -7d, -7f. The five most abundant miRs detected in ammonia stressed exosomes included members from the let7 family, let-7b, -7c, -7d, as well as miR-23a and miR-92a, both which were found in high abundance in standard cultures. Table 2 details the ten most abundant miRs in cells, MPs, exosomes among all cultures and the deposited dataset includes a comprehensive list of miRs identified from RNA sequencing. Although not as extensively studied, ammonia stress also activates p53 in cells (F. G. Wang et al., 2018). Since many stresses activate p53 (Olejniczak et al., 2018), the data may suggest that the enrichment in let-7 family miRs is characteristic of culture stress, and perhaps let-7 miR PCR assays may be used as a prognosticator of ensuing stress.

**Figure 4:**
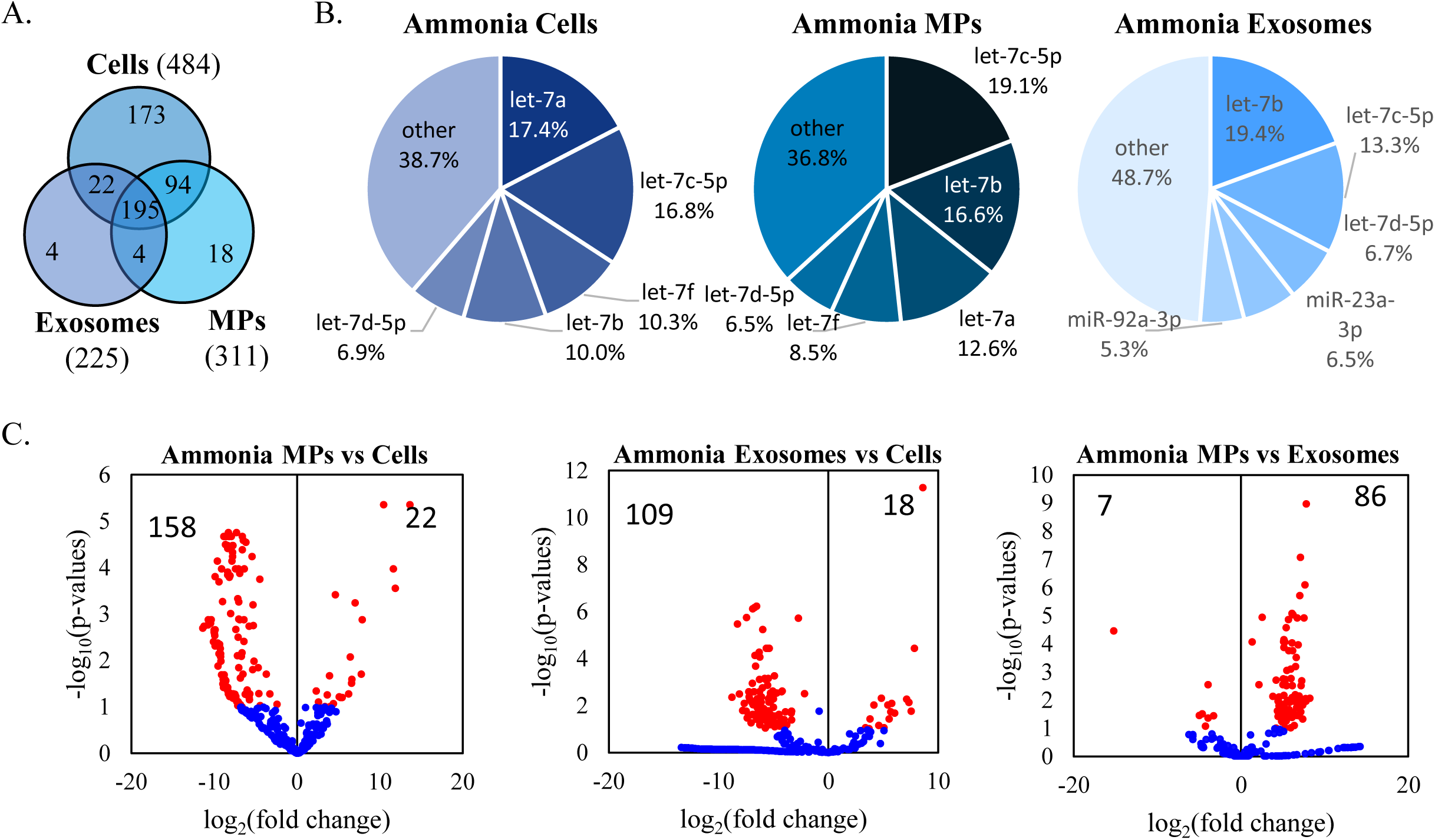
The microRNome of ammonia stressed cultures. A. Venn diagram detailing the number of unique and shared miRs amongst cells, MPs, and exosomes from ammonia stressed cultures. B. Pie chart of the relative abundance of the five most abundant miRs in cells, MPs, and exosomes from ammonia stressed cultures. The five most abundant miRs account for over 50% of the total miRs detected. C. Volcano plots of the differential expression analysis of miRs with significance determined by log_2_(fold change)>1 or <1 and -log_10_(FDR p-value)<0.1. MiR profiles are the average of three biological replicates.

Like in the osmotically stressed cultures, in the ammonia stressed cultures, the specific miR profile of the MPs most similarly reflects the miR profile of the parent cells, while the exosomes contain several of the highly enriched miRs found in the parent cells along with miRs highly enriched in standard cultures. Differential abundance levels of ammonia stressed cells, MPs, and exosomes were asymmetrical (Figure 4c). There were 158 more abundant and 22 less abuandant miRs in the ammonia stressed cells compared to the MPs. Comparing ammonia stressed cells to exosomes, there are 109 more abundant and 18 less abundant miRs. Gene ontology of the predicted mRNA targets of the ten most abundant miRs is described in supplementary table 2.

### 3.5 The shift in microRNA profiles due to ammonia and osmotic stress

The distribution of unique and differentially abundant miRs (Figure 5) and the abundance of specific miRs (Figure 6) was also compared for cell, MP, and exosome populations from the standard, ammonia stressed, and osmotic stressed culture conditions. The majority of miRs detected in cells (359 miRs) from all three culture conditions were shared (Figure 5a). Cells from standard cultures had the highest number of unique miRs (142 miRs), followed by osmotically stressed cells, 83 miRs, and ammonia stressed cells, 30 miRs. The miR profile of ammonia and osmotic stressed cells illustrates compositional differences amongst stressed conditions, where 92% (446/484 miRs) of the sequenced miRs in ammonia stressed cells were also found in osmotic stressed cells. However, 74% (396/534 miRs) of the sequenced miRs in osmotically stressed cells were found in ammonia stressed cells. Among the two stress conditions investigated, there are distinct miR profiles for each stress, illustrating that different stresses result in different miR profiles, reflecting different mechanisms of damaging of the cellular machinery and different responses to different stressors.

**Figure 5:**
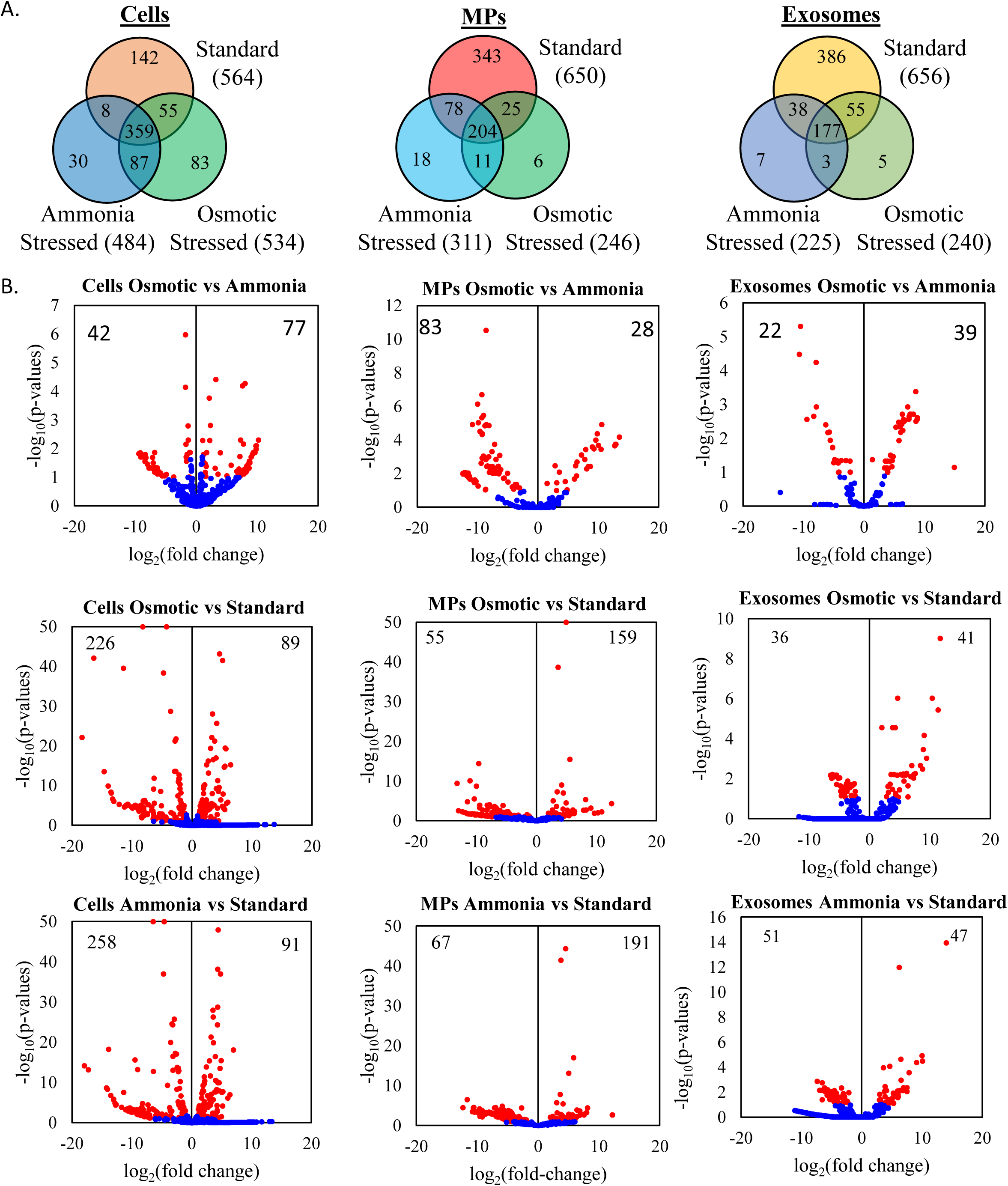
Comparing miR profile between standard, osmotic, and ammonia stressed cultures. A. Venn diagram detailing the number of unique and shared miRs of standard, osmotic, and ammonia cells, MPs, and exosomes. B. Volcano plots of the differential expression analysis of cells, MPs, and exosomes between standard, osmotic, and ammonia stressed cultures. MiRs with significance determined by log_2_(fold change)>1 or <1 and -log_10_(FDR p-value)<0.1. MiR profiles are the average of three biological replicates.

**Figure 6:**
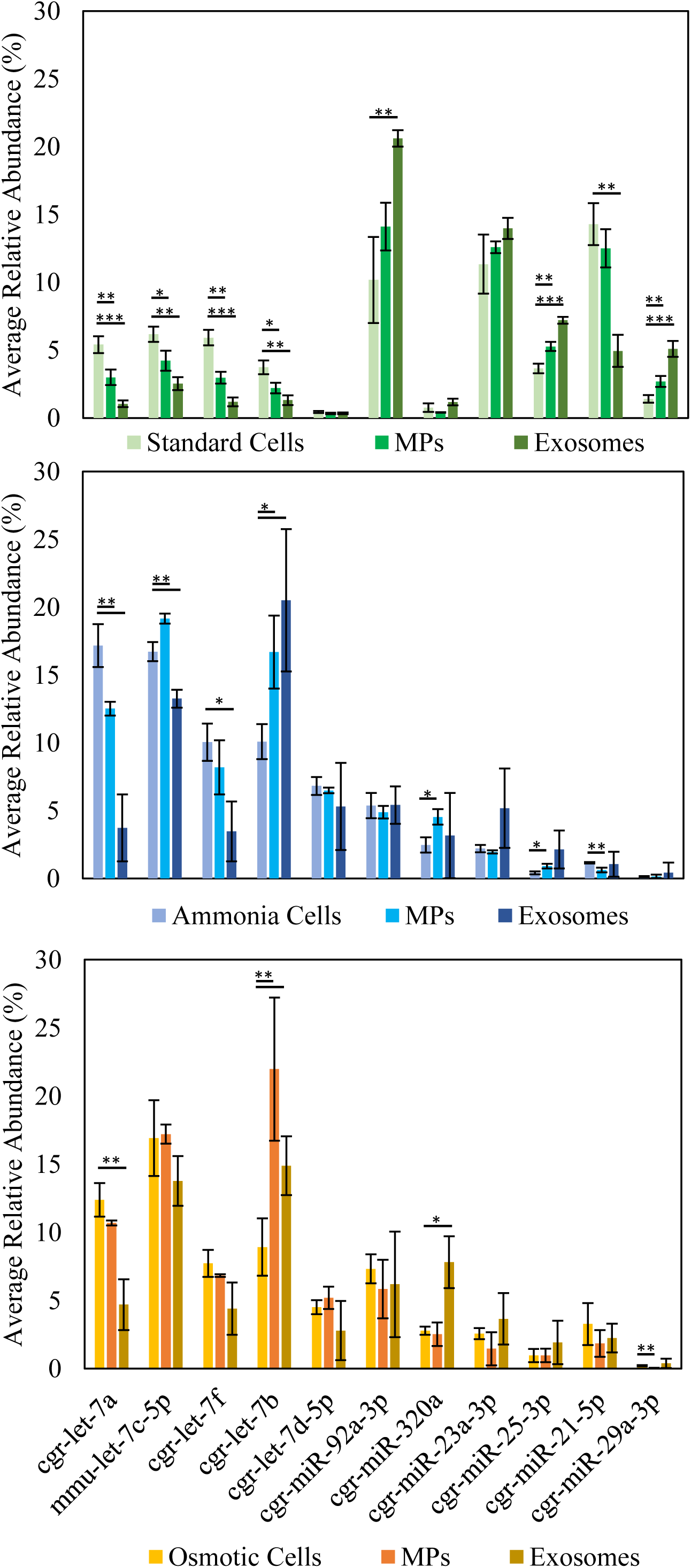
MicroRNA profile of cells, MPs, and exosomes from standard, osmotic, and ammonia stressed cultures. The relative abundance of the five most abundant miRs in each culture, standard (green), ammonia (blue), and osmotic (orange), compared between cells, MPs, and exosomes. There are significant differences in the abundance of specific miRs among cells, MPs, and exosomes of the same culture conditions, e.g., abundance of let-7a in cells, MPs, and exosomes of standard cultures are significantly different. Additionally, there is a distinct shift in miR profiles between standard cultures and stressed cultures. Three biological replicates, student t-test with p-values of * 0.05, **0.01, and ***<0.001.

The miR profiles of MPs and exosomes follow a similar trend as the cells, where MPs and exosomes from standard cultures contain a large amount of unique miRs compared to ammonia and osmotic stressed cultures. The composition of osmotically stressed MPs is 87% (215/246 miRs) similar to ammonia stressed MPs, whereas ammonia stressed MPs share 69% (215/311 miRs) of its detected miRs with osmotically stressed MPs. 75% (180/240 miRs) of the miR profile in osmotically stressed exosomes is shared with ammonia stressed exosomes and 80% (180/225 miRs) of miRs in ammonia stressed exosomes are common with osmotically stressed exosomes.

The differential miR abundances in cells, MPs, and exosomes between ammonia stress and standard culture and between osmotic stress and standard cultures display similar trends (Figure 5b) where the number of more abundant miRs in the standard cells and MPs are two to three times higher than the number of lower in abundance miRs. Differential abundance among exosomes exhibits an approximately even distribution of more and less abundant miRs.

The abundance of select, highly enriched miRs between cells, MPs, and exosomes of the same culture conditions and the between stressed and standard cultures is shown in Figure 6.

There is a distinct shift in the miR profile of stressed cultures compared to standard cultures where the let7 family of miRs was highly abundant in stressed cultures. The let7 family of miRs is present in lower relative abundance in standard cultures and conversely, highly abundant miRs in standard cultures, such as miR-92a, miR-23a, and miR-21, are present in lower relative abundance in stressed cultures. Additionally, there are significant differences in in the abundances of specific miRs between the cells, MPs, and exosomes of the same culture conditions, further illustrating that the miR composition of EVs are not simply proportional to the miR composition of the parent cell. Further, there are significant differences in specific miRs between MPs and exosomes of the same culture conditions, for example, the relative abundance of let-7a in osmotically and ammonia stressed cultures is significantly different between cells and MPs, cells and exosomes, and MPs and exosomes.

Table 3 lists the top most differentially expressed miRs in cells when comparing the two stress conditions to standard cultures, and among them. These lists have immediate functional significance regarding the role of miRs when the cells respond and adapt – as is the case here– to the two stresses: differential expression may reflect a defense response against stress or be the result of damage to cellular machinery affecting signaling and aberrant transcriptional regulation. Among the miRs in Table 3, under both stresses, notable is the consistent profound upregulation of miR-29b-5p (discussed above) miR-93-3p, and miR-15b-3p, and the consistent strong downregulation miR-706, - 1244, and -10527-5p. miR-29b-5p was discusses above and in Table 1. In human cells, miR-93 promotes cell-cycle arrest (Bao et al., 2020) by directly targeting two core cell-cycle oncogene regulators (E2F1, and Cyclin D, CCND1). In this context, miR-93 upregulation under stress is meant to inhibit proliferation in response to stress (which is consistent with the data of Figure 1), and it can thus be viewed as an indicator of stress response. miR-15 targets the anti-apoptotic BCL2 to induce apoptosis (Cimmino et al., 2005). Mir-706 is induced by ER (Wang, Han, Hu, Guo, & Chen, 2020) and oxidative stress (Yin, Guo, Zhang, & Zhang, 2016), with complex implications regarding cell survival and death. Very little is known regarding the functional role of miR-1244, and miR-10527.

**Table 3.**
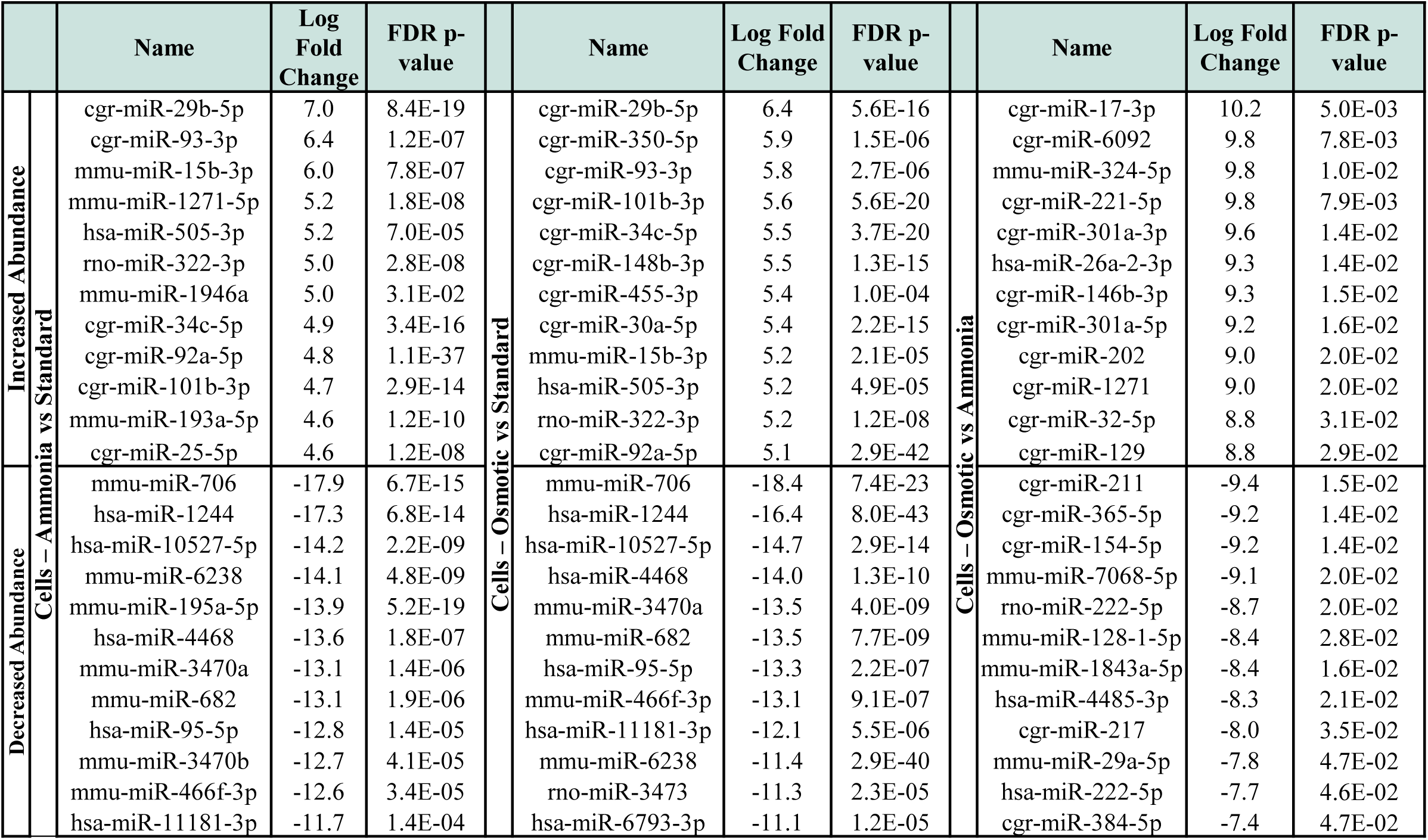

## 4. CONCLUSIONS AND A PATH FORWARD

We had previously shown that CHO cells exchange massively cytoplasmic material through EVs (Belliveau & Papoutsakis, 2022), thus suggesting that EVs have a homogenizing effect of cellular content, and notably of small non-coding RNAs, including miRs, in which EVs are highly enriched in. miRs regulate many cellular programs and phenotypes, including cell growth, metabolism, protein expression, and secretion and thus culture performance. We thus examined the miR content of cells and their small (exosomes) and large (microparticles) EVs aiming to identity the most abundant miRs under standard culture conditions and under ammonia and osmotic stress. We expect that the most abundant miRs in EVs and/or cells play a role in cell proliferation and metabolism, under standard/normal and stress culture conditions. Changes in the most abundant miRs in EVs and/or cells would reflect the impact of these miRs on cell growth, viability and metabolism.

We found that ammonia and osmotic stress alter the surface morphology of the CHO cells by abolishing the abundant surface microvilli of cells under normal culture conditions. As microvilli are likely important in the ability of cells to capture and integrate EVs, the implication is that under stress conditions there is reduced exchange of cellular material through EVs.

Under all standard and stress culture conditions, five miRs make up about 50% of the total miRs in cells, MPs, and exosomes, but these five miRs are different between standard and stressed cells and EVs. There are also substantial differences in miR content between cells and EVs and between the two types of EVs, thus demonstrating that there is an active sorting mechanism of miRs into EVs.

The two stresses result in a dramatic reduction of the total number of assayable miRs in EVs, and for the case of ammonia also in cells, thus demonstrating that the machinery that sorts miRs into EVs is damaged or inhibited by the two stresses.

The highly enriched miR-21-5p, -23a-3p and -92a-3p in standard cells and EVs (but also to a lesser extent in their stressed counterparts) are positive regulators of cell cycle and proliferation and involved in the regulation of apoptosis. All three miRs are under the control of p53, the tumor suppressor protein and transcription factor known as the guardian of the genome controlling directly or indirectly virtually all cell proliferation and cell death activities. As such, these three miRs would be targets to explore alone and in combination in cell engineering applications, and in fact, miR-92a-3p has already been explored in this context.

Under stress conditions, there is an enrichment in let-7 family miRs, occupying the top 4-5 positions in abundance in stressed cells. These miRs are known to regulate cell proliferation in a negative way and cell death in a positive way, but their impact could be more complex in the context of the imposed stressed conditions. There is clearly a need to understand their function and also if they are more broadly enriched under other physiological and genotoxic stresses. The let-7 family of miRs may be also explored as a diagnostic tool of impeding stress.

The list of highly differentially up-or downregulated miRs in cells under stress (Table 3) offers precise suggestions to explore the role of these highly differentially expressed miRs in the stress response for both diagnostic and synthetic applications.

More broadly, further studies would benefit from more extensive methods for identifying the role and specific gene targets in CHO cells of the miRs we identified to build a better understanding of the purpose of these miRs when loaded in EVs; it would also be worthwhile to determine whether these miRNAs depend on one another (joint/dual action) to enact particular phenotypes.

## Acknowledgements

We thank Deborah Powell of the Delaware Biotechnology Institute’s Bio-Imaging Center for assistance with scanning electron microscopy. Microscopy equipment was acquired with a shared instrumentation grant (S10 OD025165) and access was supported by grants from the NIH-NIGMS (P20 GM103446), the NIGMS (P20 GM139760) and the State of Delaware. We thank Bruce Kingham and Mark Shaw in the University of Delaware DNA Sequencing & Genotyping Center for assistance with the RNA library preparation and sequencing. We thank Dr. Shawn Polson and Jaysheel Bhavsar from the University of Delaware University of Delaware Center for Bioinformatics and Computational Biology Core Facility for RNAseq analysis and use of the BIOMIX compute cluster was made possible through funding from Delaware INBRE (NIGMS P20GM103446), the State of Delaware, and the Delaware Biotechnology Institute. We thank the Advanced Mammalian Biomanufacturing Innovation Center (AMBIC, NSF Grant1624698), MilliporeSigma, and Sartorius for funding. We also financial acknowledge support from Merck Sharp & Dohme Corp.

## Authorship Contributions

E.T.P. and J.B. designed the study and analyzed the data; J.B. carried out the experiments. E.T.P. and J.B. wrote the manuscript.

## Data Availability Statement

The data used to support the findings of this study are available from the corresponding author upon request. The data will be deposited in NCBI GEO in two weeks.

