## Supplemental Table 1 for "The microRNomes of Chinese Hamster Ovary (CHO) cells and their extracellular vesicles, and how they respond to osmotic and ammonia stress"

**Short Title:** CHO miRNomes & Stress

Jessica Belliveau[1,2] and Eleftherios T. Papoutsakis[1,2,3]

[1]Department of Chemical and Biomolecular Engineering, [2]Delaware Biotechnology Institute, and [3]Department of Biological Sciences, University of Delaware, Newark, DE 19711

**Corresponding Author:** Eleftherios Terry Papoutsakis

**Address:** 590 Avenue 1743, Newark, DE 19713

Papoutsakis ORCHID ID: 0000-0002-1077-1277

### Supplemental Table 1

| Comparison | Abundance | Name | Log Fold Change | FDR p-value |
|---|---|---|---|---|
| MPs – Ammonia vs Standard | Increased Abundance | mmu-miR-6395 | 12.2 | 2.3E-03 |
| | | rno-miR-298-3p | 12.2 | 2.2E-03 |
| | | mmu-miR-296-3p | 8.1 | 3.8E-05 |
| | | cgr-miR-1306-3p | 8.0 | 5.4E-04 |
| | | hsa-miR-505-3p | 7.6 | 1.4E-03 |
| | | cgr-miR-125b-3p | 7.5 | 2.1E-03 |
| | | hsa-miR-3168 | 6.9 | 2.1E-04 |
| | | rno-miR-1306-3p | 6.8 | 2.4E-03 |
| | | cgr-miR-221-3p | 6.4 | 1.2E-04 |
| | | mmu-miR-6715-3p | 6.4 | 5.3E-03 |
| | | cgr-miR-101b-3p | 6.2 | 1.6E-02 |
| | | cgr-miR-296 | 6.1 | 7.6E-03 |
| | Decreased Abundance | mmu-miR-195a-5p | -12.4 | 3.2E-05 |
| | | cgr-miR-322-5p | -11.7 | 3.2E-07 |
| | | cgr-miR-29b-3p | -10.8 | 3.1E-04 |
| | | mmu-miR-706 | -10.7 | 3.2E-04 |
| | | cgr-miR-18a-5p | -10.6 | 3.5E-05 |
| | | cgr-miR-369-3p | -10.5 | 2.9E-05 |
| | | cgr-miR-194 | -10.4 | 4.1E-04 |
| | | mmu-miR-15a-5p | -10.2 | 3.8E-04 |
| | | mmu-miR-499-5p | -9.9 | 3.7E-04 |
| | | rno-miR-15a-5p | -9.8 | 6.2E-04 |
| | | mmu-miR-330-5p | -9.8 | 7.7E-04 |
| | | cgr-miR-425-5p | -9.6 | 6.4E-05 |

| Comparison | Abundance | Name | Log Fold Change | FDR p-value |
|---|---|---|---|---|
| MPs –Osmotic vs Standard | Increased Abundance | cgr-miR-350-5p | 12.5 | 4.3E-05 |
| | | mmu-miR-1971 | 10.9 | 5.0E-03 |
| | | hsa-miR-1321 | 10.3 | 8.6E-03 |
| | | mmu-miR-28c | 9.7 | 1.1E-02 |
| | | cgr-miR-455-3p | 9.1 | 6.3E-04 |
| | | cgr-miR-34b-5p | 8.3 | 1.9E-03 |
| | | hsa-miR-3168 | 8.2 | 4.3E-06 |
| | | mmu-miR-872-3p | 7.9 | 2.0E-03 |
| | | cgr-miR-34c-5p | 6.4 | 4.0E-03 |
| | | mmu-miR-149-5p | 6.3 | 8.6E-03 |
| | | mmu-miR-378a-3p | 6.3 | 1.3E-02 |
| | | hsa-miR-25-5p | 6.1 | 1.9E-03 |
| | Decreased Abundance | cgr-miR-22-3p | -13.2 | 3.5E-10 |
| | | cgr-miR-322-5p | -13.0 | 2.7E-03 |
| | | mmu-miR-195a-5p | -12.3 | 5.6E-03 |
| | | cgr-miR-369-3p | -11.9 | 8.1E-03 |
| | | hsa-miR-7-5p | -11.6 | 1.4E-02 |
| | | cgr-miR-374-5p | -11.5 | 1.6E-05 |
| | | cgr-miR-664-3p | -11.3 | 1.8E-02 |
| | | cgr-miR-365-3p | -11.1 | 8.3E-11 |
| | | mmu-miR-350-3p | -10.9 | 2.1E-02 |
| | | cgr-miR-29b-3p | -10.8 | 1.5E-02 |
| | | mmu-miR-706 | -10.7 | 1.6E-02 |
| | | mmu-miR-203-3p | -10.6 | 1.7E-02 |

| Comparison | Abundance | Name | Log Fold Change | FDR p-value |
|---|---|---|---|---|
| MPs – Osmotic vs Ammonia | Increased Abundance | cgr-miR-34b-5p | 13.5 | 6.5E-05 |
| | | cgr-miR-361 | 12.9 | 1.7E-04 |
| | | cgr-miR-455-3p | 12.8 | 1.7E-04 |
| | | mmu-miR-98-3p | 12.7 | 2.2E-04 |
| | | mmu-miR-149-5p | 10.5 | 1.2E-05 |
| | | cgr-miR-350-5p | 10.4 | 3.4E-04 |
| | | mmu-miR-872-3p | 10.0 | 2.0E-04 |
| | | cgr-miR-425-5p | 9.8 | 4.1E-05 |
| | | mmu-miR-7044-5p | 9.6 | 9.9E-05 |
| | | hsa-miR-4492 | 9.2 | 3.3E-04 |
| | | mmu-let-7b-3p | 9.0 | 2.5E-04 |
| | | hsa-miR-1321 | 8.7 | 3.7E-03 |
| | Decreased Abundance | mmu-miR-18a-3p | -12.4 | 9.3E-03 |
| | | cgr-miR-30e-3p | -12.4 | 1.1E-02 |
| | | mmu-miR-29b-1-5p | -12.3 | 9.5E-03 |
| | | cgr-miR-15b-3p | -12.2 | 8.9E-03 |
| | | rno-miR-222-3p | -12.1 | 8.2E-03 |
| | | cgr-miR-31-5p | -11.9 | 1.2E-02 |
| | | cgr-miR-125b-3p | -11.7 | 1.3E-02 |
| | | mmu-miR-671-5p | -11.6 | 1.6E-02 |
| | | mmu-miR-296-3p | -11.5 | 9.2E-03 |
| | | mmu-miR-1271-5p | -11.5 | 1.3E-02 |
| | | cgr-miR-324-3p | -11.4 | 1.3E-02 |
| | | rno-miR-1306-3p | -11.4 | 1.5E-02 |

| Comparison | Abundance | Name | Log Fold Change | FDR p-value |
|---|---|---|---|---|
| Exosomes – Ammonia vs Standard | Increased Abundance | hsa-miR-3168 | 14.1 | 1.2E-14 |
| | | hsa-miR-1321 | 10.1 | 3.4E-05 |
| | | hsa-miR-4796-3p | 10.0 | 1.2E-05 |
| | | cgr-miR-574 | 9.1 | 4.3E-05 |
| | | mmu-miR-674-5p | 7.9 | 2.8E-04 |
| | | hsa-miR-6131 | 7.5 | 4.6E-03 |
| | | hsa-miR-12135 | 7.5 | 9.0E-03 |
| | | cgr-miR-148b-3p | 6.9 | 4.6E-03 |
| | | hsa-miR-4502 | 6.9 | 7.4E-03 |
| | | rno-miR-450b-5p | 6.9 | 5.3E-03 |
| | | hsa-miR-505-3p | 6.8 | 7.5E-03 |
| | | mmu-miR-6239 | 6.5 | 1.1E-03 |
| | Decreased Abundance | cgr-miR-365-3p | -7.4 | 1.4E-03 |
| | | cgr-miR-186-5p | -7.0 | 7.4E-03 |
| | | mmu-miR-100-5p | -6.6 | 7.3E-03 |
| | | cgr-miR-409-3p | -6.6 | 4.1E-02 |
| | | cgr-miR-425-5p | -6.5 | 1.8E-03 |
| | | mmu-miR-99a-5p | -6.1 | 4.7E-03 |
| | | cgr-miR-20a | -6.1 | 4.7E-03 |
| | | cgr-miR-151-3p | -6.1 | 8.9E-03 |
| | | cgr-miR-17-5p | -5.9 | 7.5E-03 |
| | | cgr-miR-664-3p | -5.8 | 9.3E-03 |
| | | rno-miR-409a-3p | -5.7 | 1.8E-02 |
| | | mmu-miR-92a-3p | -5.6 | 6.8E-03 |

| Comparison | Abundance | Name | Log Fold Change | FDR p-value |
|---|---|---|---|---|
| Exosomes – Osmotic vs Standard | Increased Abundance | hsa-miR-3168 | 11.7 | 9.3E-10 |
| | | cgr-miR-486-3p | 11.4 | 3.6E-06 |
| | | cgr-miR-672 | 10.4 | 9.3E-07 |
| | | rno-miR-1896 | 9.5 | 9.2E-04 |
| | | cgr-miR-140-3p | 9.0 | 6.8E-05 |
| | | cgr-miR-450b-3p | 8.9 | 3.5E-04 |
| | | mmu-miR-5106 | 8.8 | 3.3E-03 |
| | | hsa-miR-1321 | 8.4 | 2.2E-03 |
| | | mmu-miR-378a-3p | 7.6 | 5.4E-03 |
| | | mmu-miR-5099 | 7.3 | 6.0E-03 |
| | | hsa-miR-4796-3p | 7.3 | 9.1E-03 |
| | | mmu-miR-674-5p | 7.0 | 2.2E-03 |
| | Decreased Abundance | cgr-miR-369-3p | -6.5 | 6.5E-03 |
| | | mmu-miR-100-5p | -6.3 | 9.4E-03 |
| | | cgr-miR-365-3p | -6.2 | 6.0E-03 |
| | | cgr-miR-34c-3p | -5.9 | 8.0E-03 |
| | | mmu-miR-99a-5p | -5.8 | 6.5E-03 |
| | | cgr-miR-425-5p | -5.4 | 6.5E-03 |
| | | cgr-miR-664-3p | -5.4 | 1.9E-02 |
| | | mmu-miR-350-3p | -5.2 | 9.4E-03 |
| | | cgr-miR-27b-3p | -5.0 | 8.7E-03 |
| | | cgr-miR-99a-5p | -4.8 | 2.3E-02 |
| | | cgr-miR-374-5p | -4.8 | 4.2E-02 |
| | | cgr-miR-139-5p | -4.7 | 7.0E-02 |

| Comparison | Abundance | Name | Log Fold Change | FDR p-value |
|---|---|---|---|---|
| Exosomes – Osmtic vs Ammonia | Increased Abundance | cgr-miR-450b-3p | 14.9 | 7.1E-02 |
| | | cgr-miR-486-3p | 8.8 | 2.4E-03 |
| | | cgr-miR-409-3p | 8.8 | 2.7E-03 |
| | | cgr-miR-672 | 8.6 | 3.1E-03 |
| | | cgr-miR-340-5p | 8.5 | 4.1E-04 |
| | | hsa-miR-34b-3p | 8.2 | 1.9E-03 |
| | | cgr-miR-186-5p | 7.7 | 1.9E-03 |
| | | cgr-miR-140-3p | 7.6 | 1.9E-03 |
| | | rno-miR-214-3p | 7.5 | 2.2E-03 |
| | | mmu-miR-709 | 7.4 | 2.7E-03 |
| | | cgr-miR-28-3p | 7.2 | 1.2E-03 |
| | | cgr-miR-652-3p | 6.8 | 2.9E-03 |
| | Decreased Abundance | rno-miR-702-3p | -10.7 | 3.3E-05 |
| | | cgr-miR-574 | -10.5 | 4.9E-06 |
| | | cgr-miR-1839-5p | -9.4 | 2.7E-03 |
| | | hsa-miR-1260b | -8.3 | 2.2E-03 |
| | | cgr-miR-151-5p | -7.9 | 5.6E-05 |
| | | cgr-miR-148b-3p | -7.9 | 1.2E-03 |
| | | mmu-miR-1983 | -6.3 | 3.9E-03 |
| | | cgr-miR-486-5p | -6.0 | 6.6E-03 |
| | | mmu-miR-10a-5p | -5.7 | 6.4E-03 |
| | | cgr-miR-139-5p | -5.6 | 1.1E-02 |
| | | cgr-miR-328 | -5.2 | 1.8E-02 |
| | | cgr-miR-15b-5p | -5.2 | 1.8E-02 |

### Supplemental Table 2

| GO Term – Molecular Function Standard Cells | P-Value | FDR |
|---|---|---|
| RNA polymerase II core promoter proximal region sequence-specific DNA binding | 1.2E-08 | 9.9E-06 |
| Transcriptional activator activity, RNA polymerase II transcription regulatory region sequence-specific binding | 7.4E-05 | 0.028 |
| RNA polymerase II transcription factor activity, sequence-specific DNA binding | 0.00012 | 0.028 |
| Transcription factor activity, sequence-specific DNA binding | 0.00013 | 0.028 |
| Sequence-specific DNA binding | 0.00031 | 0.053 |

| GO Term – Molecular Function Standard MPs | P-Value | FDR |
|---|---|---|
| RNA polymerase II core promoter proximal region sequence-specific DNA binding | 7.30E-07 | 0.00056 |
| Transcriptional activator activity, RNA polymerase II transcription regulatory region sequence-specific binding | 0.00023 | 0.0794 |
| Ubiquitin protein ligase binding | 0.00031 | 0.079 |

| GO Term – Molecular Function Standard Exosomes | P-Value | FDR |
|---|---|---|
| RNA polymerase II core promoter proximal region sequence-specific DNA binding | 3.5E-06 | 0.0029 |
| DNA binding | 4.6E-05 | 0.019 |
| Ubiquitin-protein transferase activity | 0.00027 | 0.056 |
| Transcriptional activator activity, RNA polymerase II transcription regulatory region sequence-specific binding | 0.00027 | 0.056 |
| ATP binding | 0.00053 | 0.087 |
| SMAD binding | 0.00063 | 0.087 |

| GO Term – Molecular Function Osmotic Cells | P-Value | FDR |
|---|---|---|
| RNA polymerase II core promoter proximal region sequence-specific DNA binding | 9.0E-06 | 0.0062 |
| SMAD binding | 0.00014 | 0.049 |
| MAP kinase tyrosine/serine/threonine phosphatase activity | 0.00040 | 0.072 |
| Chromatin binding | 0.00056 | 0.072 |
| Zinc ion binding | 0.00063 | 0.072 |
| RNA polymerase II transcription factor activity, sequence-specific DNA binding | 0.00073 | 0.072 |
| Ubiquitin protein ligase activity | 0.00073 | 0.072 |
| DNA binding | 0.0011 | 0.087 |
| Protein dimerization activity | 0.0011 | 0.087 |

| GO Term – Molecular Function Osmotic MPs | P-Value | FDR |
|---|---|---|
| RNA polymerase II core promoter proximal region sequence-specific DNA binding | 6.8E-05 | 0.041 |

| GO Term – Molecular Function Osmotic Exosomes | P-Value | FDR |
|---|---|---|
| RNA polymerase II core promoter proximal region sequence-specific DNA binding | 1.9E-06 | 0.0014 |
| Chromatin binding | 2.1E-05 | 0.0079 |
| Kinase activity | 0.00031 | 0.076 |
| Transcriptional activator activity, RNA polymerase II transcription regulatory region sequence-specific binding | 0.00057 | 0.092 |
| GDP binding | 0.00062 | 0.092 |

| GO Term – Molecular Function Ammonia Cells | P-Value | FDR |
|---|---|---|
| RNA polymerase II core promoter proximal region sequence-specific DNA binding | 4.6E-05 | 0.032 |

| GO Term – Molecular Function Ammonia MPs | P-Value | FDR |
|---|---|---|
| RNA polymerase II core promoter proximal region sequence-specific DNA binding | 2.4E-05 | 0.016 |

| GO Term – Molecular Function Ammonia Exosomes | P-Value | FDR |
|---|---|---|
| RNA polymerase II core promoter proximal region sequence-specific DNA binding | 5.3E-05 | 0.035 |
